# A ready-to use workflow for automated, sound-based bird identification and localization

**DOI:** 10.1101/2024.07.02.601711

**Authors:** Carsten M. Buchmann, Frank M. Schurr

**Affiliations:** Institute of Landscape and Plant Ecology, University of Hohenheim, Ottilie-Zeller-Weg 2, 70599 Stuttgart, Germany

**Keywords:** Passive acoustic monitoring, bird species identification, automated workflow, Audiomoth, BIRDnet, localization

## Abstract

Field studies of bird communities typically require the fine-scale mapping of individuals. Passive acoustic monitoring combined with localization of individuals is a promising approach to gather such data. While various approaches for identification of species and localization of individuals have been proposed, no fully automated ready-to-use workflow is available so far. We here present a novel approach based on sound recordings with multiple cost-efficient automated recording units (Audiomoths). The workflow uses a well-established AI model (BirdNET; other models possible) for species identification and localizes the sources of all identified bird sounds with high accuracy. Tests with replayed sounds of different bird species in an agricultural landscape show that - after filtering out identifications with low identification confidence - the algorithm localizes more than 90% of the sounds within 5 m of the true location (85 % < 2 m). Recording and localization of wild birds demonstrate the applicability of the approach for avian ecology. This workflow is completely automated and ready-to-use, also for non-experts and can also be used when strong winds affect the speed of sound or if 3D localizations are of interest. By making data on individual bird locations accessible the presented work will help to advance fundamental and applied ecology as well as conservation.

## Introduction

Localizing organisms in space is central to ecology (Krebs 1972). In avian ecology, it is essential for bird censuses and for quantifying behaviour and habitat use. There is thus a strong need for simple, cost- and labour-efficient approaches to localize birds.

The efficiency of bird monitoring has increased massively through the widespread use of passive acoustic monitoring (PAM) with automated recording units (ARU; e.g. Darras, Batáry et al., 2018, Ruff et al., 2020 and references therein for examples of different taxa, Pérez-Granados & Traba, 2021).

Recently, low-cost ARUs for deployment in the wild have become available (Smith et al., 2022, Manzano-Rubio et al., 2022). Among these the Audiomoth by Open Acoustic Devices (Hill et al., 2019) is to our knowledge the most widely used model nowadays since it provides good performance in relation to the low price of below 100 $ (Darras et al., 2019, Lapp et al., 2023).

The combination of audio recordings of wildlife with artificial intelligence, e.g. convolutional neural networks (CNNs) for automated species identification, is a rapidly developing research area (e.g. Ruff et al., 2020, Manzano-Rubio et al., 2022, Müller et al., 2023). Among numerous models for automated species identification (seed Xie et al., 2023 for a comprehensive review) for birds the CNN BirdNET (Kahl et al., 2021) currently seems to be the state-of-the-art tool since it is open source and comparatively simple and ready-to use. It was evaluated as the method of choice, particularly if one is interested in a large array of species and has limited knowledge in bird identification based on sounds, and hence cannot easily provide a large set of training data needed for more specific CNN approaches (Lauha et al., 2022, Michaud et al., 2023, Xie et al., 2023).

A next step besides pure species identification is the localization of the detected individuals. Pérez-Granados and Traba (2021) give a comprehensive overview of methods to use ARUs for density estimation. They assess the use of microphone arrays (i.e. several synchronized microphones distributed in space) that enable the localization and distinction of different individuals based on time delays of sounds on different recordings as most promising (see Rhinehart et al., 2020 for a summary on time delay-based methods). However, high costs and logistic effort as well as time required for the interpretation of the recordings are so far major disadvantages of this method (Pérez-Granados & Traba, 2021).

In recent years, various studies used microphone arrays to localize individuals of single species (Wilson & Bayne, 2018, Avots et al., 2022) or limited numbers of species (Smith et al., 2022). They used expensive technology (beyond 1000 $ per ARU; Avots et al., 2022) in combination with manual species identification (Smith et al., 2022), and laborious manual detection of time delays (Wilson & Bayne, 2018, Smith et al., 2022). Bistel et al. (2022) present a set of algorithms including complex CNNs that can do both identification and localization and tested it on synthetic sounds of one species in a small-scale lab-setting (<15 m). This approach is, however, far from an automated ready-to-use workflow, for instance because the AI models still needs to be trained for every case study and there is no simple application or running code available that can be used directly for collected field data. What is still missing, but urgently needed, is a cheap and simple, ready-to-use integrated workflow. Moreover, such a workflow should be able to identify a large number of species and localize individuals in a fully automated manner.

Here, we present a workflow that uses low-cost sound recorders (Audiomoths, Hill et al., 2019) and an existing AI for bird species identification (BirdNET, Kahl et al. 2021), to calculate accurate positions of individual birds – all entirely automated. This workflow thus offers a cost-efficient way to identify and localize individual birds in the field. We test the approach in both a simple and a structurally complex landscape.

## Materials and Methods

### Use of ARUs and setup in the field

For localization of the origin of sounds based on time delays, the sound needs to be detected by at least three microphones (ARUs). The detection distance of the microphones should be considered for the spatial setup of the ARUs. The locations of the ARUs need to be recorded with high accuracy (e.g. high precision differential GPS). The ARUs should be set up to start at the same time to simplify temporal calibration (synchronization), see below. If the duration of time gaps between predefined recording periods on different ARUs cannot be guaranteed to be consistent between different ARUs, it is recommended to plan the observation period within a non-interrupted recording session (one WAV file per ARU).

We used cost-efficient Audiomoth ARUs (Model 1.2.0, Open Acoustic Devices, Hill et al., 2019) in this study. In a preliminary test for these devices we had found a good detection distance of bird sounds up to 100m facing the sound and 50-100 m facing the opposite direction, dependent on background noise, e.g. wind. Gaps between recording periods as well as recording speed are not consistent to an acceptable degree (between five tested Audiomoths we had found differences of up to 1.3 s per file gap when recording continuously and up to 0.03 s runtime differences of 60 minute recordings). For this reason, the recording period was set to fall in a non-interrupted recording session and a temporal calibration with two calibration signals was performed.

After deployment and (simultaneous) start of all ARUs in the study site, a loud sound signal for temporal calibration/synchronization that could be recorded by all ARUs was emitted from a defined position (loud clapping). Alternatively, an external signal (e.g. church bells from a known position) can also be used if it can be recognized and temporally located on all recordings with high precision. At the end of the observation period, another calibration signal was given. These signals were used to correct for small differences in starting times and recording speeds of individual ARUs.

### Temporal calibration of ARU recordings (Fig. 1.1)

We used Audacity (free software), to extract the timing of the calibration signals on each ARU recording. The sounds can be visualized, the signals marked and the markers exported as a so-called label file (TXT). The accuracy that can and should ideally be reached with this approach is < 0.003 s (corresponding to a sound travel distance of 1 m). From the times of the calibration signals on each recording, any event on any recording can be calibrated to the average time of all recordings by linear interpolation. The time needed for the sound to travel from the calibration point (where the calibration signal is emitted) to any ARU was considered in the calculation. These calibrated times of any recorded sound denote the standardized time since the first calibration signal.

**Fig.1.**
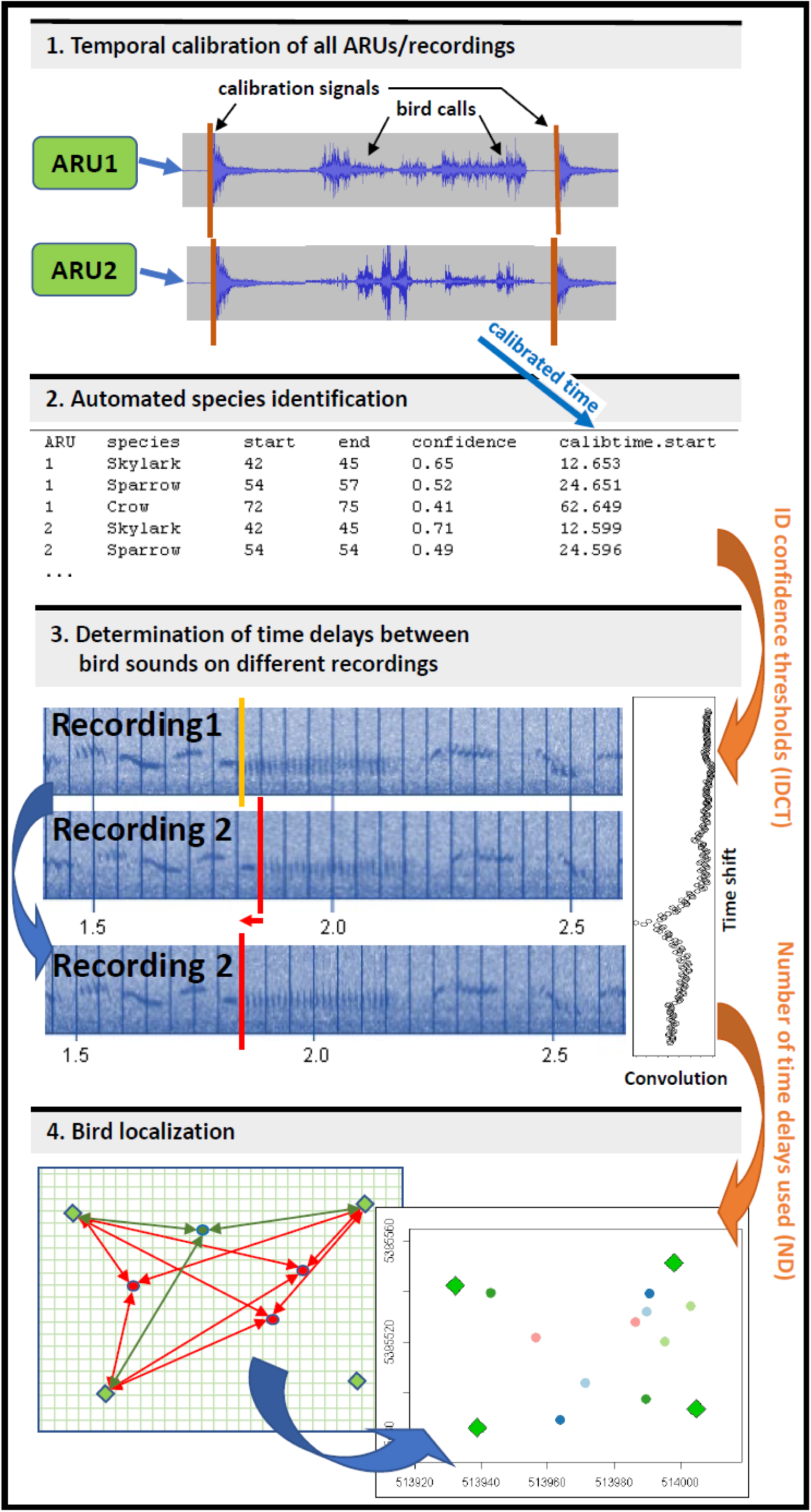
An automated workflow of bird identification and localization. Red arrows indicate parameters that were varied in this study. 1.: The exact times of the calibration signals are identified on all recordings. 2.: Sets of identifications belonging to the same bird sound are identified from the output of the ID algorithm (here BirdNET), these sets are filtered by confidence thresholds and the calibrated time for each identification is calculated. 3.: Time delays between the same sound signal on different recordings (sound file snippets of 3 seconds) are determined by determining the time shift yielding the highest spectrogram similarity by means of convolution. 4.: For a grid of potential sound locations (red points as examples) surrounding the sample area, the RMSE between the expected and observed time delays is calculated. This procedure is then repeated around the grid point with smallest RMSE (green point) using a finer grid.

### Species identification (Fig. 1.2)

All recordings were analysed with BirdNET-Analyzer (Kahl et al., 2021, Software Version 2022), which produces a file that lists all species identified including the confidence of the identification in three second time slots (e.g. 6-9, 12-15, 33-36 s etc.) for the entire recording (from now on referred to as ‘ID hits’). Other methods for the identification can be used, if the time slot and identification confidence of any sound event are provided. We recommend time intervals of 3 s (as is the default in BirdNET) to cover large parts of a bird song or call but to still reduce the chance of a mixture of several sounds of different individuals or species within time intervals to a large extent. Besides using AI species identification this task can also be by experts (in the field or listening to the recordings), in which case the (real) times of the individual observations need to be provided for further analysis.

### Analysis and determination of delays of sound signals (Fig. 1.3)

Sets of ID hits (‘ID sets’) were identified that were found on at least three recordings, i.e. the same species in a 3 s time slot with a calibrated start time of the time slot that is not more different from the others than a threshold (here set to 0.75 s; note: by starting the recordings at the same time, the time slots are largely synchronous so that this threshold should not become relevant). Subsequently a two-step filter was applied: first, only ID hits exceeding a certain confidence threshold were considered (parameter IDCT 1, here set to 0.25), and secondly at least one of the ID hits from the ID set needed to exceed another confidence level (parameter IDCT 2, here set to 0.75, also tested: 0.35). Then a sound snippet of the corresponding 3 s interval was created (software R, package TuneR, R Core Team, 2021, Ligges et al., 2023) for all recordings and the calibrated time of the start of each snippet was stored with it.

For the automated determination of time delays of a sound signal on different recordings, the different snippets were converted into spectrograms (R package signal, Signal developers, 2023). Frequencies below 1000 and above 20000 Hz were excluded eliminating background noise but still retaining the relevant frequencies for bird sounds. Subsequently the phase information was discarded, spectrograms were converted to dB, scaled and centred. Then these spectrograms were compared with respect to similarity (see below). This comparison was done between one reference recording (the recording where the ID hit had the highest ID confidence) and each of the other recordings.

To calculate time delays between sound signals, the spectrogram of each recording was shifted relative to the spectrogram of the reference recording (pixel by pixel; for a default recording setting of 48kHz a pixel represents 0.0027 s, which is a good compromise between runtime and accuracy). The temporal overlap for shifting the 3 s spectrograms relative to each other was set to 2.7 s. This means that in the beginning of the shifting procedure 0.3 s at the end of the reference recording and at the beginning of any other recording are not considered and vice versa at the end of the procedure (i.e. only 2.7 s are compared). Shifting the spectrograms by up to 0.3 s can detect delays for a situation where the distance of two ARUs to a sound source is different by up to ∼ 100 m. At each shifting step, similarity of the spectrograms was measured as the convolution of the two sound signals (the sum of all pairwise dB products for corresponding time and frequency). The time shift at which the similarity between the compared spectrograms is highest defines the time delay of the sound signal relative to the reference. The relative difference between maximum similarity and average similarity across all time shifts considered was recorded as a general performance measure indicating the quality of the time delay assessment, and could be used to filter the results later on. Together with the calibrated time of each snippet start this gives the total time delay between detections of a sound signal by any pair of ARUs. As alternative to the automated approach, time delays can also be determined manually/optically (e.g. in Audacity) and provided for further analysis.

### Localization of sound sources (Fig. 1.4)

Knowing at least two time delays of a sound signal (from three recordings) allows the localization of the sound origin. One can decide to use only the best ones with respect to the performance measure (see above), or more, up to the maximum number that are available for a specific sound (parameter ‘ND’ for number of time delays used, here set to 3, also tested: maximum available). For any point in a coarse 2D grid (we here used 5 m by 5 m, 100 m around the sample area) the corresponding time delays were calculated based on the speed of sound from this point to the locations of the ARUs (2D distance; note: wind speed can be considered here, see below). These expected time delays were compared to the measured time delays by means of RMSE. This procedure was then repeated for a finer 2D grid (here 0.5 m by 0.5 m) covering 10 m x 10 m around the best location found using the coarse grid. Subsequently, 3 D localization was performed by sampling a cube (0.5 m by 0.5 m by 0.5 m, 10 m around the best point), considering 3D distances and, if applicable, 3D wind velocities from potential sound locations to the ARUs, to assess the height of the sound origin.

The sonic speed (according to conditions, notably temperature) and wind speed as well as direction during the observation period can be specified by the user and are then considered by the algorithm. Wind is specified as a 3D velocity vector with x pointing East, y pointing North and z pointing upwards (we here used 335 m s^-1^ and (0,0,0) m s^-1^, for a recording at 7°C without wind). The relevance of incorporating wind for accurate localization of sound sources can be illustrated with a small example: in a situation of wind 20 m s^-1^ from the West, and a sound location in between two ARUs at 50 m distance to each, all aligned with the wind, the signal arrival times at the ARUs correspond to those expected from a sound location is shifted approx. 3 m to the West without wind. Our algorithm corrects for this bias by calculating any relevant speed of sound by vector addition of the respective sound velocity vector and the wind velocity vector.

### Field studies to test the identification and localization workflow

In an agricultural field site (Agricultural Research Station Heidfeldhof of the University of Hohenheim, Stuttgart, Germany) we set up six Audiomoth ARUs (Model 1.2.0, Hill et al., 2019) in a rectangle of approx. 70 m by 60 m (Fig. 3) in March 2023. In this study sound recordings were done to develop and test the algorithm and to evaluate different parameter settings (‘validation study/site’). Audiomoth ARUs were programmed to start at the same time, recording continuously at 48kHz with medium gain and mounted on metal poles at 1.5m height. We emitted a calibration signal from another defined position in the centre of the area by clapping two small metal boxes 9 times. From six defined positions, 1.5 m above ground, we then played bird sounds of five different species (Carrion Crow, Corvus corone, file: XC758247, White Wagtail, *Motacilla alba*, file: XC712328, Eurasian Skylark, *Alauda arvensis*, file: XC717648, Eurasian Tree Sparrow, *Passer montanus* , file: XC737729, Meadow Pipit, *Anthus pratensis*, file: XC760273; https://xeno-canto.org/). Each bird sound was played 13 times over the course of approx. 30 minutes with a smartphone and a bluetooth speaker (JBL Flip Essential) at 75% of maximum volume in six different defined locations distributed in the rectangle of ARUs. After that a second calibration signal was emitted and the recordings were stopped. All positions were measured with a Trimble GeoX DGPS (differential correction in postprocessing resulted in an accuracy of 78% < 0.15 m, 90% < 0.3 m, 99% < 1 m). Wind was < 2 m s^-1^ without predominant direction during the study, hence neglected in the analysis.

A second set of recordings were collected in a traditional orchard with larger trees adjacent to a forest (at University Hohenheim Research Station Unterer Lindenhof, Ehingen, Germany) to apply the approach in a more closed landscape (‘application study/site’) in April 2023. Here we again used six Audiomoth ARUs arranged in a rectangle of approx. 100 by 60 m(Figs 6a,b). To test the algorithm in an environment with strong wild bird activity, the same bird sounds were again played with the speaker, now 18 times in six defined locations. In this study, however, the total recording period was approx. 1.7 h to also record and subsequently analyse sounds of wild birds. In this study the spatial accuracy of the defined locations was: 40% < 0.15 m, 69% < 0.3 m, 93% < 1 m. Wind speed and direction were again neglectable in this study.

## Results

The localization algorithm was able to locate the origin of bird sounds (handheld speaker) that were identified by the AI (here: BirdNET) with moderate confidence levels (thresholds: 2 or more recordings > 0.25, one of these > 0.75) for most species with high accuracy. While for Carrion Crow no set of recordings passed these thresholds and for Eurasian Tree Sparrow only approx. 10%, for Eurasian Skylark, White Wagtail and Meadow Pipit all sounds passed the ID confidence thresholds and > 90% of sounds were located with an accuracy of less than 5 m, >85% of less than 2m, > 50% of less than 1m (Figs 2, 3).

**Fig. 2.:**
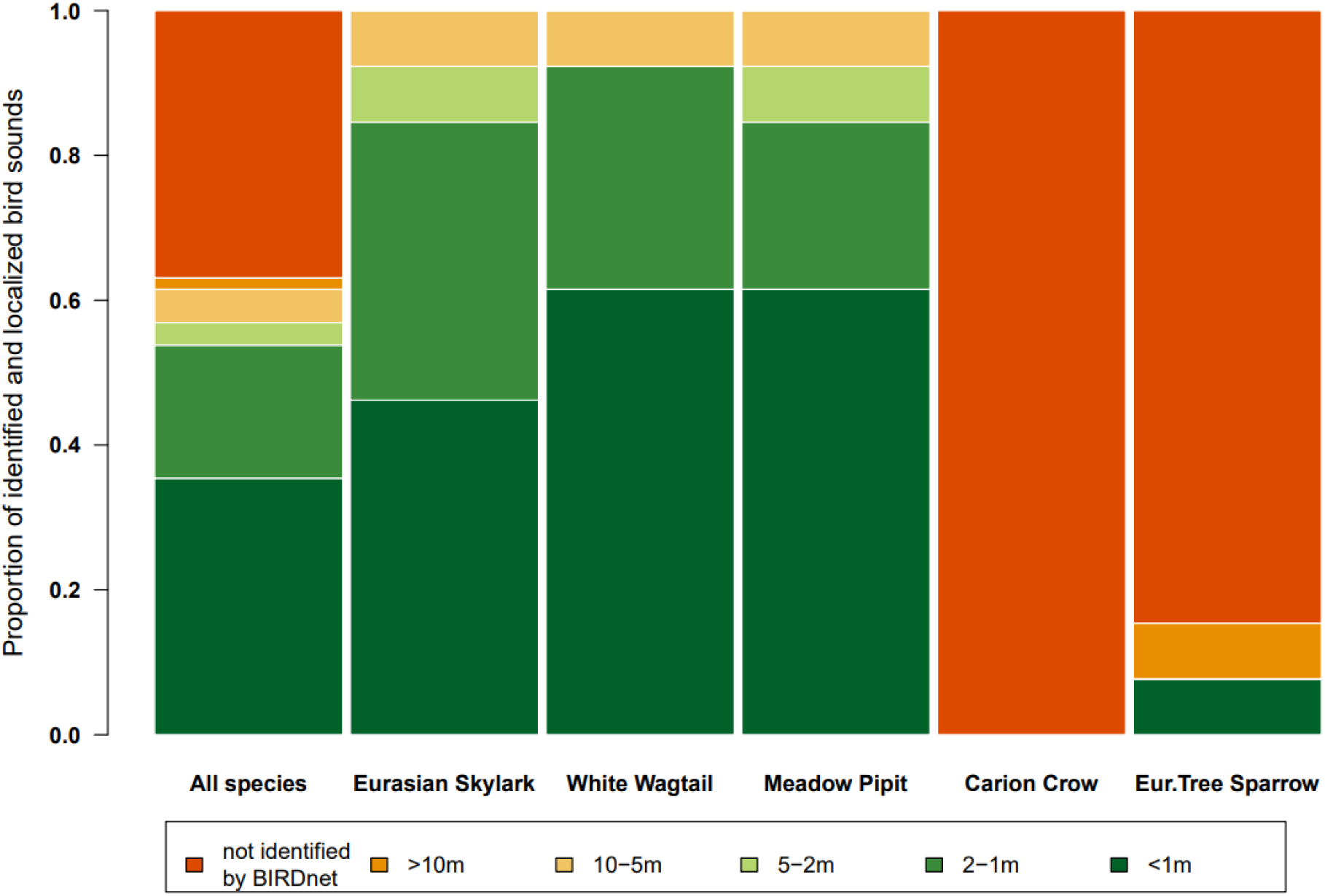
Proportion of sounds of different bird species replayed in the validation experiment that were correctly identified by BirdNET and localized with a certain accuracy.

**Fig. 3:**
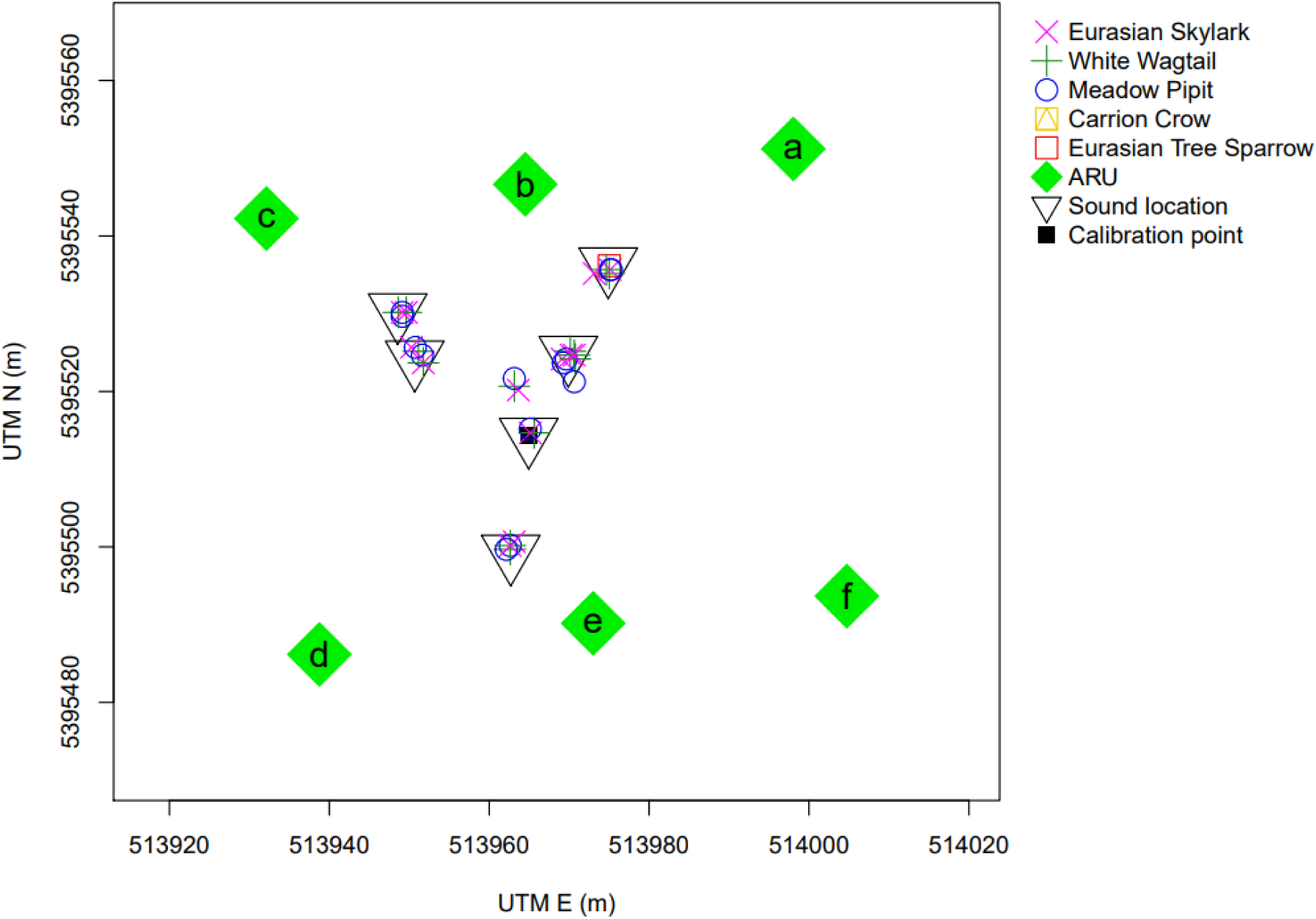
Setup of the validation experiment in an agricultural landscape, showing the locations of ARUs, the calibration point (calibration signal emitted), the locations of bird sound playback and the localization of these bird sounds by the algorithm (UTM Zone 32U).

The presented localization algorithm calculates 2D and 3D positions of the sound source. Since our validation site was flat and ARUs as well as the playback locations of bird sounds were at the same height, considering the 3rd dimension did not impact our results for the 2D locations of sound origins. The resulting z-coordinates were, however, randomly distributed in the scope of values that were given to be tested by the algorithm (20 m). This is not surprising, since ARUs mounted in a plain can gather little information on different heights of a sound source.

Two parameters controlling the localization algorithm were found to affect localization accuracy namely the second ID confidence threshold (IDCT2) and the number of time delays used for localization (ND). We found that a higher IDCT2, while lowering the number of sound events that enter the actual localization, increases the accuracy of these localizations (Fig. 4). If a sound is on n recordings n-1 time delays are availble; the quality of their determination is quantified with a performance measure based on the success of the spectrogram overlay anaylsis using convolution, compare Fig 1.3). Localization accuracy increased further if only the three most clearly defined time delays (ND) were used instead of using all time delays that were available for any sound (Fig. 4).

**Fig. 4.**
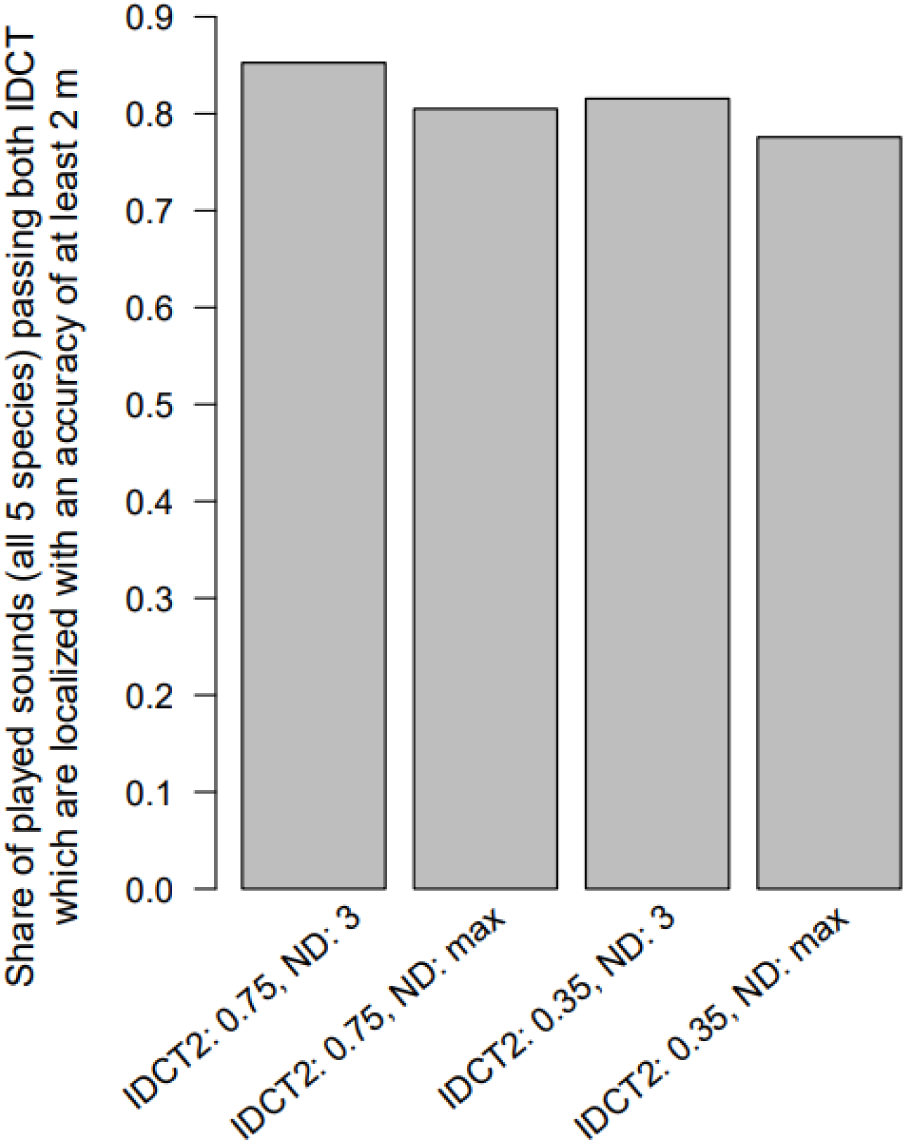
Effects of the second ID confidence threshold (IDCT2)and number of time delays used for localization (ND) on the proportion of correctly identified bird sounds that were localized within 2 m of the true location. Note: 100% here denotes the number of sounds that passed IDCT1 and IDCT2.

To test how inaccuracies in quantifying delay times affect localization accuracy, we added random noise (n = 250, normal distribution, mean = 0, SD =0.0025 s resulting in 95% of values within 0.01 s around the true value; this is oriented towards the fine vertical elements in the spectrogram of a Eurasian Skylark sound, compare Fig. 1) to calculated delay times for different sound locations, and used these to re-estimate the sound location (Fig. 5). In the area delimited by the ARUs, 95% of localizations were < 2m from the original sound location. This localization accuracy decreased as sound locations moved away from this area (Fig. 5).

**Fig. 5:**
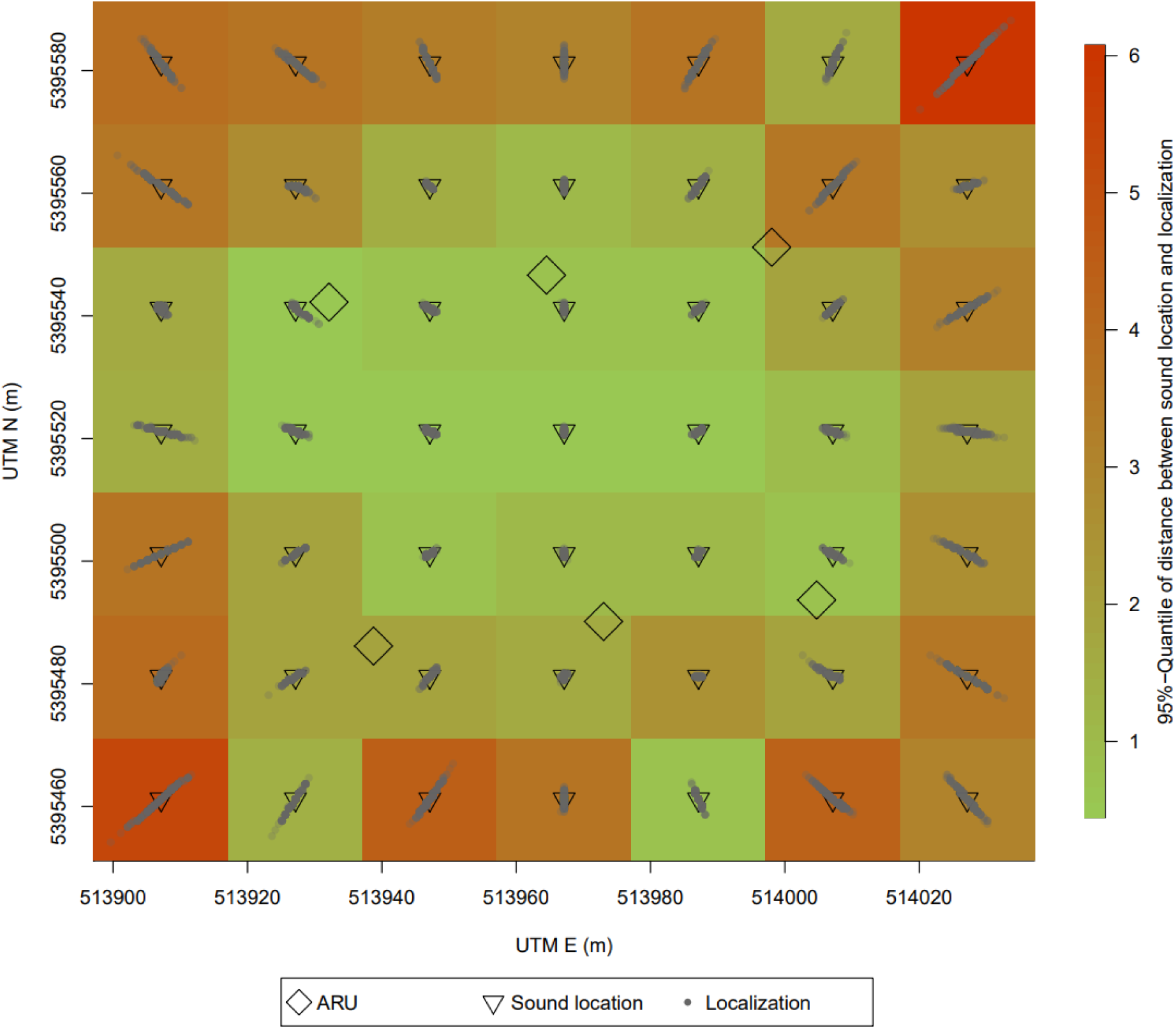
Error propagation from inaccuracies in the determination of time delays to the localization of sound origins in the validation experiment (UTM Zone 32U).

In the application study in a more closed landscape (orchard next to a forest) where many birds where actively singing, a lower number of replayed bird sounds passed the ID thresholds (Fig. 6a). However, of these sounds 93 % were localized with an accuracy of < 5 m (61 % < 2 m). In the application study we also localized all natural bird sounds that BirdNET identified in the 1.7 h recording period. This yielded 202 localizations of 9 bird species (Fig. 6b). These can be used to derive habitat utility descriptions for different species or even individuals (based on the known time for each localization), such as the estimation of home ranges (for an example see Fig. 6b).

**Fig. 6:**
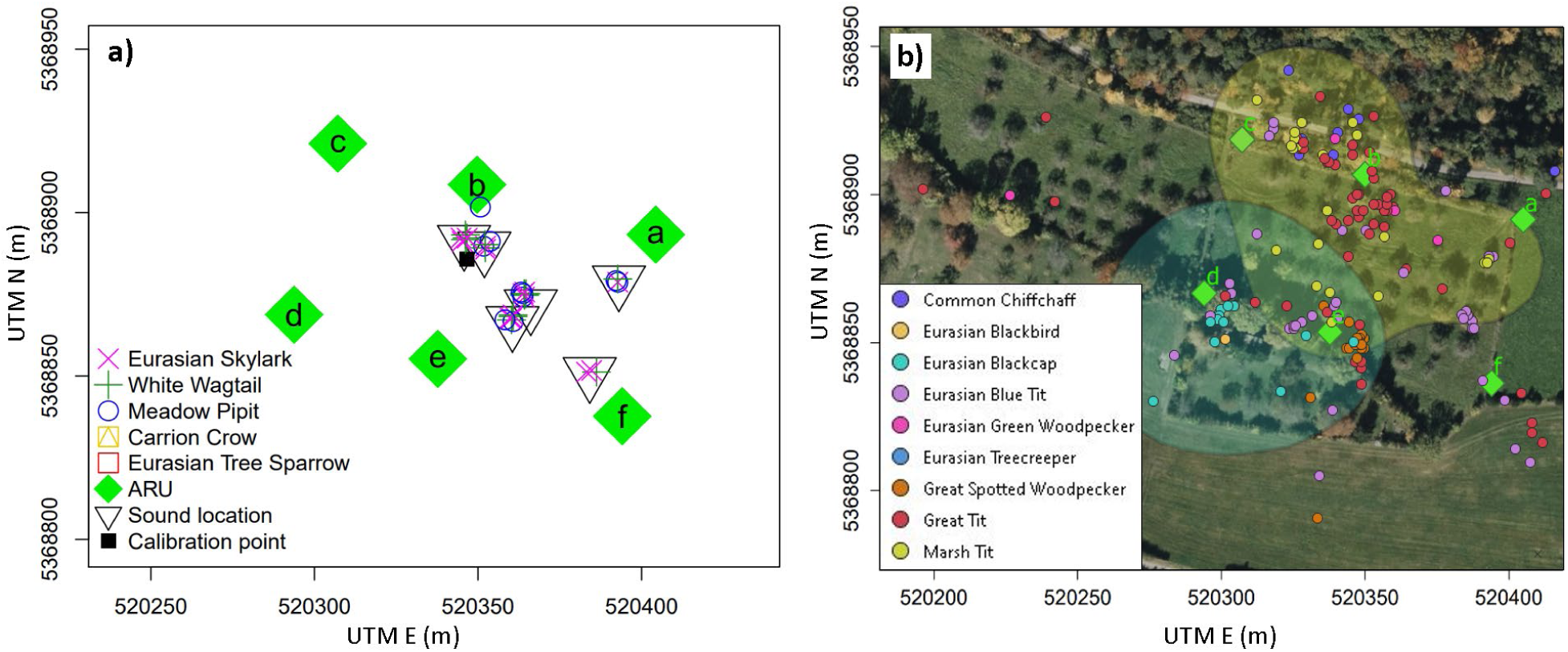
Localization result of the played bird sounds (a) during the application study at the natural field site (orchard close to forest), and bird localizations of bird sounds beyond the manually played ones during the recording period lasting for 1.7 h (b). In (b) the polygons illustrate the 75 % utility density based on the localizations of two species, Marsh Tit and Eurasian Blackcap (function getverticeshr in R package adehabitatHR) as a possible application of localization results (background photo from Bingmaps).

## Discussion

The presented automated workflow can facilitate bird monitoring, conservation and fundamental ecological research by identifying and precisely locating birds based on their sounds. It requires a simple setup in the field and uses comparatively low-cost sound recorders. The localization analysis is coded in a ready-to-use R script which can be executed easily without advanced programming skills. If required, the workflow also enables localization in 3D space and can correct for bias imposed by strong winds.

In recent years other studies have presented solutions for different aspects of our workflow. While BirdNET currently seems to be the AI of choice for multi-species systems (compare Toenies & Rich, 2021, Cole et al., 2022, Singer et al., 2024, for a comparison to other methods see Xie et al., 2023), the development of AI systems for species identification can be expected to be fast in the next years. Any novel output from possibly more advanced AI models in the future can be easily incorporated into our workflow without much adjustment. Some recent studies have used time delays on different recordings for localization of wildlife. However, these have either only done so for single to few species, used comparatively expensive technology or require manual determination of time delays. Hence, these approaches are still far from ready-to use for multi-species analyses (e.g. Bistel et al. 2022, Smith et al., 2022, Barré et al., 2024). While our workflow still allows for both alternatives, manual identification of sounds by experts and manual/visual detection of time delays, the complete analysis presented here, using AI for species identification and the automated time delay assessment and subsequent localization, is the first cost-efficient, simple and ready-to-use workflow for identification and localization individual birds in species-rich systems. Still, AI for identification provided, the approach can also be applied to other taxa that can be detected by sound.

### The value of automated bird localization

The ongoing decline in bird diversity globally (Rosenberg et al., 2019, Burns et al., 2021), and in European farmland birds specifically (Traba & Morales, 2019, Rigal et al., 2023), calls for efficient monitoring to quantify trends for different species and understand the underlying drivers. Passive acoustic monitoring (PAM) has considerable potential in this respect but faces the unresolved challenge of how to convert the number of recorded bird calls into an estimate of population density (Pérez-Granados & Traba, 2021). Knowing the exact location and time of any bird call overcomes this challenge by making it possible to apply standard methods of distance sampling and home range mapping (Bibby et al. 2000; Buckland 2006).

Considering that habitat destruction and loss of landscape heterogeneity is a major driver of declining bird abundance (Xu et al., 2018, Traba & Morales, 2019, Rigal et al., 2023) understanding fine scale habitat use of individuals is crucial for targeted conservation efforts (e.g. Blackburn & Cresswell 2015). Here a method that combines low-disturbance PAM with accurate localization of individuals can significantly improve our understanding of fine-scale resource and habitat use of species and individuals including movement behaviour and how these are affected by landscape changes. Also, fundamental ecological research on biotic interactions in natural communities as well as the assessment of ecosystem services such as seed dispersal (Schurr et al. 2009) or pollination (Schmid et al., 2015) by birds, can substantially benefit from small-scale spatial information of bird activity and movement.

### Challenges, limitations and extensions of the presented approach

The temporal calibration of ARUs is a simple procedure that allows the use of comparatively cheap equipment, but can cause some disturbance for the birds. If available, an external sound signal (like bells from a close-by church tower) can be used to avoid disturbance. If a more advanced ARU system is available, where the start and end times of recordings are synchronous (<0.002 s, e.g. through cable or network connection) and recording speed is comparable, or GPS-equipped ARUs (e.g. GPS board for Audiomoths) can be used, which provide a comparable time stamp for all recordings, the temporal calibration procedure can be omitted. In this case an interrupted recording scheme (e.g. 15 minutes once every hour) can also be used, since calibration signals at the beginning and the end of every non-interrupted recording period would not be required.

As in all PAM methods the detection distance for the species - ARU combination of interest needs to be considered and tested before deploying the ARUs. If a larger area shall be sampled in a grid of ARUs especially the detection distance of the ARUs in all directions need to be considered for choosing a grid resolution which ensures that any sound originating from the sampling area can be identified on at least three recordings. For Audiomoths, our test indicated a detection distance that is reduced by up to ∼ 50% when not facing the sound origin. The fact that also more advanced and expensive ARU systems do not have considerably larger detection distances (compare Darras, Furnas et al., 2018, analysing detection distances, and Darras et al., 2019, comparing prices of ARU systems) points to the possibility but also the economic reasoning to use cheap ARUs, especially when larger areas are to be monitored and numerous ARUs are thus required.

In our study, we applied a general rather low threshold for identification confidence (0.25) for all recordings and a second higher threshold (0.75) for at least one identification of a set of recordings belonging to one bird sound. While the first one is simply filtering very unlikely identifications, the latter is a reasonable value for most species (see Pérez-Granados, 2023, and references therein for a more detailed discussion on confidence thresholds). Still applying these thresholds had different consequences for sounds of different species, namely that from 0 % to 100 % of sets of identifications (from different ARUs) belonging together were excluded. Within species lowering this second confidence threshold of identifications entering the localization procedure decreased the accuracy of localizations. For any real field monitoring study lowering the confidence score (i.e. accepting an identification more easily) moreover increases the risk of false identifications and hence wrong conclusions. Our results and those from several recent studies illustrate that confidence levels have to be chosen with care and possibly need to be adjusted (and validated) for different species (e.g. Barré et al., 2019, Cole et al., 2022, Metcalf et al., 2022, Singer et al., 2024). Setting different thresholds for different species could be easily incorporated in the analysis (exemplary code is included). Still, the approach to handle and deal with confidence scores is a topic that currently receives a lot of attention and can be expected to soon yield new insights that will help to improve this aspect of our workflow.

In the tested setup (ARUs at the same height) the 3D localization did not yield reasonable information on the height of the sound source. A little example can illustrate the reason: in a situation where two ARUs are 100m apart and a sound source is in between at 70/30m distance, raising the height of the sound source by 15 m changes the distances to the ARUs only slightly (1.5 and 3.6 m) and hence causes a change of the time delay between the recordings of both ARUs of only 0.0058 s, which is very small compared to the resolution of the temporal calibration and the spectrogram analysis. Hence small inaccuracies in the determined time delays (by spectrogram comparisons) cause comparatively large variation in the z coordinate that is calculated. For reasonable use of 3D localization ARUs need to be mounted at different heights, and for a specific localization of any sound only cases should be considered where the respective recordings used for this localization also stem from ARUs at different heights.

The number of time delays (pairs of ARUs) used for localization also needs to be considered. Ignoring time delays for which the time delay could not be determined precisely improved the localization performance (Fig. 4). However, while in principle enough, only using two delays (i.e. three recordings) can be problematic if all three ARUs are placed in a row (which can happen if ARUs are set up in a grid array), since then a sound origin could be located at the same distance left or right of this line. Hence, we recommended to use at least three time delays. If ARUs are set up at a distance that is oriented towards their detection distance it can almost be ruled out that for a specific sound source four ARUs in a row, and not ARUs surrounding the sound, would deliver the clearest time delays and hence would be chosen for the localization.

Since during our studies there were no strong winds from a predominant direction this aspect was not considered (wind velocity vector was set to (0,0,0) m s^-1^). Still, in situations such as a sample area on a slope in times of falling winds it can be crucial to specify all three wind velocity dimensions.

## Conclusions

This study presents the first ever ready-to-use workflow that, after a simple setup using affordable equipment, combines species identification with individual localization for birds, both fully automated. The species identification can be performed by experts or using available AI models for the organism of interest. Since this was found to be the most critical aspect for missing out bird sounds, expected future advances in the field of AI for species identification will strongly benefit our approach. The subsequent localization approach can in principle be used for any sound source. Application of the workflow in environments where 3D localization is relevant, for example aquatic systems, or in situations with strong wind, is possible without larger adjustments. Being able to identify and localize bird individuals will help to advance monitoring and develop targeted conservation measures for species. Also, fundamental ecological research on species interactions, behavioural ecology and ecosystem services and functioning can substantially benefit from a simple and fast workflow providing novel data on individual movements at high spatial and temporal resolution.

## Acknowledgements

We thank S. Klos, M. Kasten and T. Köhler for helping in the field and running preliminary studies. Moreover, we thank the teams of the University Hohenheim research stations Heidfeldhof and Unterer Lindenhof.

## Author contributions

CMB and FMS conceived the ideas for the study and the methodology; CMB lead the field study and collected the data; CMB and FMS developed the workflow, CMB analysed the results and led the writing of the manuscript. All authors contributed critically to the drafts and gave final approval for publication.

## Data availability

A zip-archive including R-code of the entire analysis together with exemplary data can be downloaded from: https://tinyurl.com/yhv4tcaj ( these will be published on figshare after publication)

## Notes

https://tinyurl.com/yhv4tcaj

### Competing Interest Statement

The authors have declared no competing interest.

